# Cellular energy regulates mRNA translation and degradation in a codon-specific manner

**DOI:** 10.1101/2023.04.06.535836

**Authors:** Pedro Tomaz da Silva, Yujie Zhang, Evangelos Theodorakis, Laura D. Martens, Vicente A. Yépez, Vicent Pelechano, Julien Gagneur

**Affiliations:** School of Computation, Information and Technology, Technical University of Munich, Garching, Germany; Munich Center for Machine Learning, Munich, Germany; Department of Microbiology, Tumor and Cell Biology, Karolinska Institutet, Stockholm, Sweden; Computational Health Center, Helmholtz Center Munich, Neuherberg, Germany; Institute of Human Genetics, School of Medicine, Technical University of Munich, Munich, Germany

## Abstract

**Background:** Codon optimality is a major determinant of mRNA translation and degradation rates. However, whether and through which mechanisms its effects are regulated remains poorly understood.

**Results:** Here we show that codon optimality associates with up to 2-fold change in mRNA stability variations between human tissues, and that its effect is attenuated in tissues with high energy metabolism and amplifies with age. Biochemical modeling and perturbation data through oxygen deprivation and ATP synthesis inhibition reveal that cellular energy variations non-uniformly affect the decoding kinetics of different codons.

**Conclusions:** This new mechanism of codon effect regulation, independent of tRNA regulation, provides a fundamental mechanistic link between cellular energy metabolism and eukaryotic gene expression.

## Background

Codons encode in three nucleotides 20 amino acids in a redundant way. Importantly, codons coding for the same amino acid, or synonymous codons, are not functionally equivalent. In particular, codons differ on their optimality for being decoded by the translation machinery, which not only affects the rate of protein production but also the rate of messenger RNA (mRNA) degradation [1], via a pathway termed codon optimality-mediated mRNA degradation [2]. Mechanistically, a ribosome dwelling at a given non-optimal codon not only delays its translation but can also be recognized by the Ccr4-Not complex triggering mRNA degradation [3].

Differences in optimality between codons have been suggested to be mostly determined by variation in cognate tRNA concentration (reviewed in [4]). In light of such an explanation, regulation of the tRNA pool composition between cell types and conditions could differentially affect mRNA translation and degradation. Consistent with this hypothesis, associations between codon usage and differential expression between tissues and conditions have been reported across eukaryotes [5–9]. However, whether regulation of the tRNA pool is generally causing these associations is debated. While variations of the tRNA pool have been reported by some studies [5,8,10], other studies, including some with advanced tRNA sequencing protocols [11–13], reported surprisingly stable proportions of the tRNA pool per anticodon. Hence, under which conditions and how codon optimality plays a role in differential gene expression remains unclear.

## Results

Here we analyzed the effect of codon optimality on differential gene expression by first looking at mRNA stability. To this end, we considered changes in the ratio between exonic and intronic RNA-Seq read coverage as a proxy for the variation of mRNA stability ([14], Methods). We processed 7,771 RNA-Seq human post-mortem samples from the GTEx project spanning 49 tissues and 528 individuals (Fig. 1A, Fig. S1). Typically, mRNA stability varied 2.7-fold between tissues (median fold change between lowest and highest decile). In comparison, mRNA abundance typically varied by 4.2-fold between tissues (transcript per million median fold-change between lowest and highest decile). This observation shows that mRNA stability variations substantially contribute to between-tissue mRNA regulation and underscores the importance of post-transcriptional processes in gene regulation, in agreement with previous studies [15,16].

**Fig. 1:**
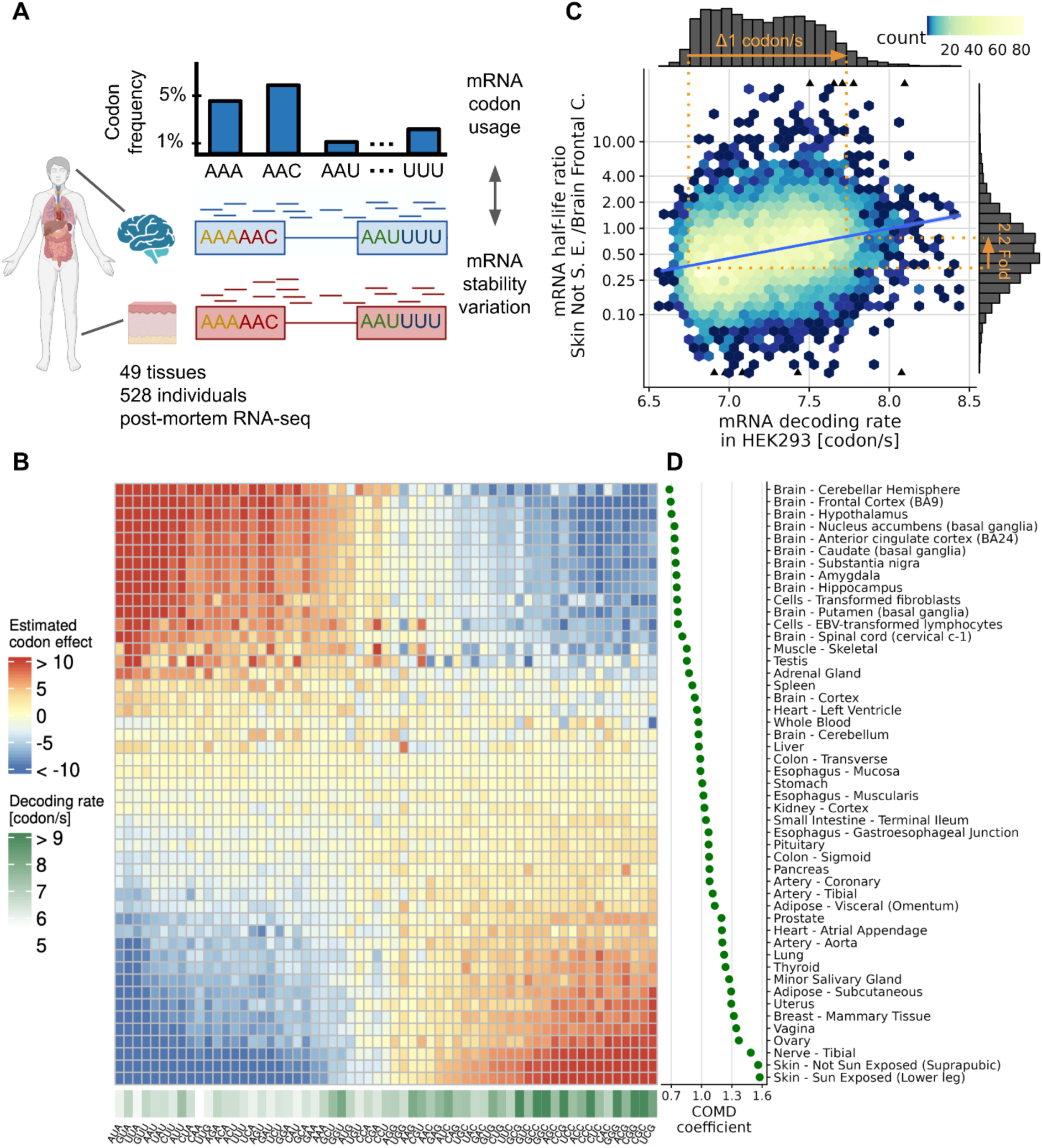
Codon optimality associates with mRNA stability variations between tissues. **A** Variations in mRNA stability for every gene are estimated using the ratio of the number of exonic reads, mostly reflecting the balance between mRNA synthesis and degradation, to the number of intronic reads, mostly reflecting mRNA synthesis [14]. We then investigated how those estimated variations in mRNA stability associate with codon usage. **B** Association (regression slope, Fig. S2) between codon frequency (column) and relative mRNA half-life for each tissue (rows), along with codon decoding rate measured in the HEK293 cell line (lower track), a measure of codon optimality. The tissues are ordered by increasing COMD coefficient (right panel, Methods). **C** mRNA half-life ratio of suprapubic skin to frontal cortex against mRNA decoding rate. The mRNA decoding rates are expressed in codons per second and computed using codon decoding rates measured in HEK293 cells (Methods). According to the linear regression (blue line), a decoding rate change of 1 codon per second in HEK293 associates with 2.2-fold larger mRNA stability ratios (orange annotations). **D** COMD coefficient across tissues.

To study mRNA stability regulation, we defined the relative half-life as the ratio of the mRNA stability in a tissue to its mean across tissues (Methods). We observed a striking association pattern between codon frequencies and tissue-specific relative half-life. Codons considered optimal according to different measures of codon optimality exhibit a distinct association with tissue-specific relative half-life compared to non-optimal codons (Fig. 1B, Fig. S3). We next asked whether these codon-level associations were also reflected at the mRNA level. To this end, we considered for each mRNA its decoding rate in HEK293 cells as a codon optimality measure [17]. The dynamic range of HEK293 mRNA decoding rates spanned about 1 codon per second (difference between the slower and the faster 4^th^ percentile, Fig. 1C). Remarkably, variation in this range was associated with changes in half-life of surprisingly large amplitudes, including a 2.2 fold-change between suprapubic skin and frontal cortex (Fig. 1C). These results suggest that the amplitude of the effect of codon optimality on mRNA degradation is modulated and is an important contributor to mRNA regulation across human tissues.

To systematically study variations of codon optimality effects on mRNA half-life, we introduced a new metric termed codon optimality-mediated degradation (COMD) coefficient. The COMD coefficient quantifies how much the relative half-life of an mRNA is predicted to change given a decoding rate increase of 1 codon per second in HEK293. The higher the COMD coefficient in a given sample, the more beneficial the usage of optimal codons for mRNA stability. The COMD coefficient was maximal in sun-exposed skin, reaching a value of 1.6 fold-change/codon/s (Fig. 1D). Assuming causality, this suggested that recoding an mRNA such that its decoding rate is 1 codon per second faster in HEK293, could increase its half-life by 1.6-fold in sun-exposed skin relative to the average tissue. The COMD coefficient was minimal in several brain tissues with values of about 0.7 fold-change/codon/s.

We next asked which biological processes, if any, could cause this apparent modulation of the effect of codon optimality on mRNA degradation. Using the Gene Ontology, we found the strongest associations for mitochondrial ATP synthesis pathways, the expression of whose genes negatively correlated with the COMD coefficient across tissues (Gene set enrichment analysis, False Discovery Rate, FDR < 10^−6^ for Mitochondrial ATP synthesis coupled electron transport and other mitochondrial ATP synthesis related GO terms, Fig. 2A, B, Table S1). Furthermore, this negative correlation was also observed across individuals within the same tissues (Fig. 2C, Fig. S4C).

**Fig. 2:**
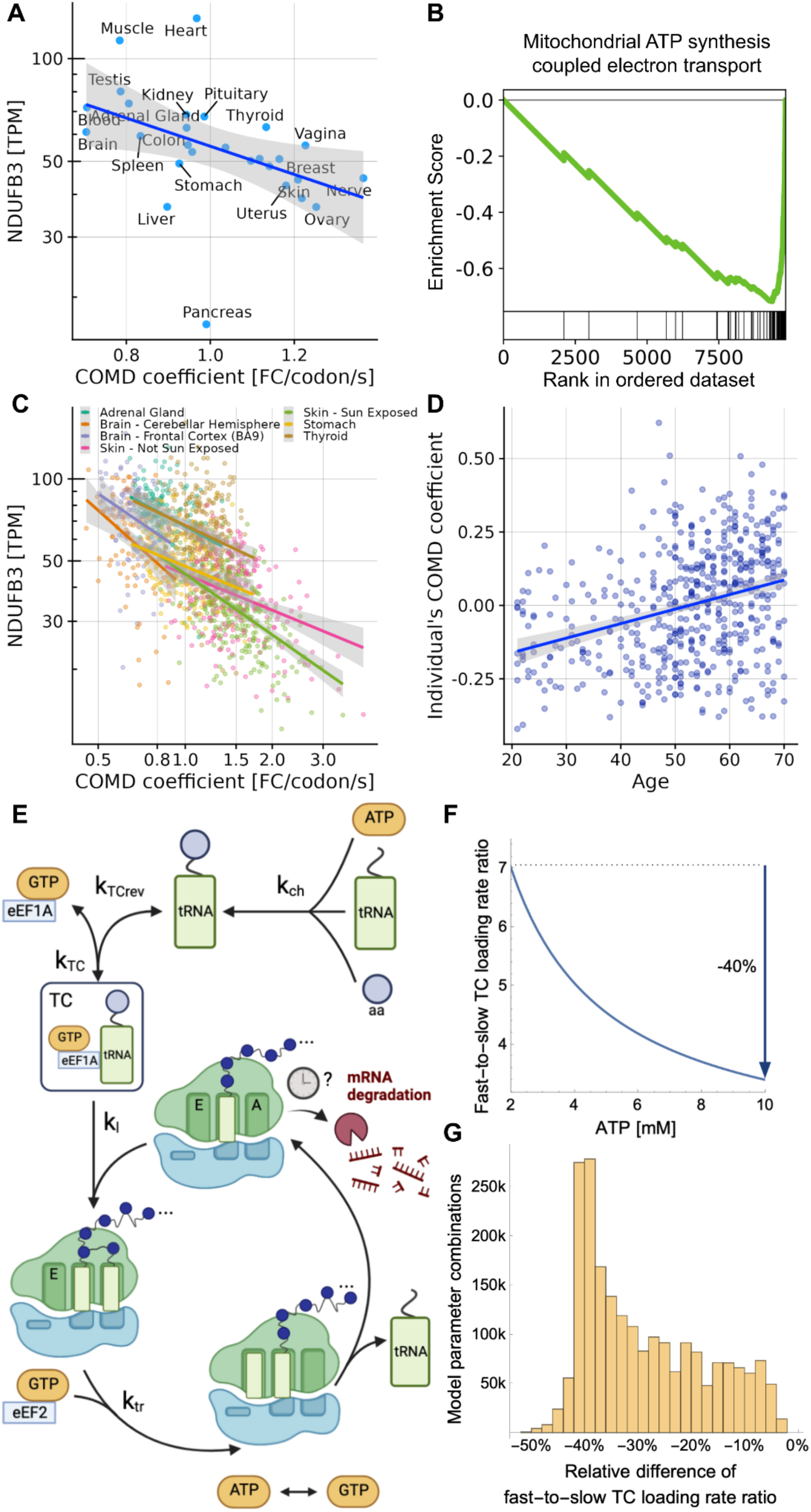
ATP is predicted to modulate relative codon decoding rates. **A** Expression in transcripts-per-million (TPM) across major GTEx tissues of *NDUFB3*, a representative nuclear-encoded gene that encodes a protein of the mitochondrial respiratory chain, against the COMD coefficient (Spearman’s rho = −0.62, *P =* 7.5×10^−4^). **B** Gene set enrichment analysis [18] for the Gene Ontology biological process “mitochondrial ATP synthesis coupled electron transport” in human. **C** *NDUFB3* expression (TPM) against the COMD coefficient across GTEx individuals for 7 tissues. **D** COMD coefficient of GTEx individuals estimated adjusting for tissue (Methods) against age (Spearman’s rho = 0.33, *P =* 9×10^−15^). **E** Reactions considered in our biochemical model of the eukaryotic translation elongation cycle (top to bottom): tRNA aminoacylation, ternary complex (TC) formation, TC loading, ribosome translocation, and ATP to GTP conversion. The mRNA degradation is depicted but not included in the model. **F** Fast-to-slow TC loading rate ratio against ATP concentration for a representative combination of plausible kinetic rate constants (Methods). The model predicts that the fast-to-slow TC loading rate ratio is 40% lower for the maximal compared to the minimal ATP concentration. **G** Distribution of the relative difference between the minimal and maximal ATP concentrations of the fast-to-slow TC loading rate ratio across combinations of plausible model parameter values. Across all parameter combinations, the TC loading rate ratio is lower for maximal ATP concentrations.

To assess the generality of these observations beyond the GTEx dataset, we next processed single-cell RNA-Seq data from 45,146 mouse cells [19] that we aggregated into 45 cell types. In mouse cells, mitochondrial ATP synthesis genes also negatively correlated with the COMD coefficient (Gene set enrichment analysis, FDR < 0.01, Fig. S4A-C). Taken together, these observations suggest that the more reduced the mitochondrial ATP synthesis, the more beneficial to mRNA stability the usage of optimal codons. Consistent with this hypothesis, we furthermore found that age, which is linked with a decline in mitochondrial function (reviewed in [20]), positively correlated with the COMD coefficient (Fig. 2D, GTEx dataset).

How could ATP mechanistically modulate how much codon optimality impacts mRNA stability? Translation is one of the most energy-demanding processes in the cell [21]. In order to decode one codon at least 3 energy-carrier molecules are needed, 1 ATP and 2 GTP [22,23]. Mitochondria produce energy in the form of ATP, which regenerates GTP [24]. We hypothesized that changes in ATP abundance, and therefore GTP as well, alter the kinetics of the translation elongation cycle unequally for different codons and consequently how likely the Ccr4-Not complex gets recruited and triggers mRNA degradation. To formally explore this hypothesis, we developed a biochemical model of the translation elongation cycle (Fig. 2E, Methods).

In a translation elongation cycle, the ternary complex (TC), which is composed of one aminoacyl-tRNA, one GTP, and one eukaryotic translation elongation factor 1A (eEF1A), is first loaded into the A-site of a translating ribosome. Next, the amino acid is added to the nascent polypeptide chain and the ribosome translocates freeing up the A-site for a new cycle [22]. If TC loading is slow, the E-site can be freed while the A-site remains empty, setting the ribosome into a conformation recognized by the Ccr4-Not complex [3]. Hence, the faster the TC gets loaded on the ribosome, the less likely mRNA degradation is triggered.

Our biochemical model predicts that higher amounts of ATP increase the fraction of tRNAs in TC and, consequently, the TC loading rate (Fig. S5). Moreover, the model describes how this relationship depends on the overall abundance of the tRNA. Non-optimal codons, for which the abundance of cognate tRNAs is low [2], show a relatively higher TC loading rate increase as ATP increases compared to optimal codons. As a result, the TC loading rate ratio of optimal to non-optimal codons decreases with ATP concentration (Fig 2F). This qualitative behavior was robust to choices of rate constants within a broad range of plausible values (Fig 2G). Altogether, these theoretical investigations support a model in which cellular energy attenuates the effects of codon optimality by dampening the impact of overall cognate tRNA abundance variation on TC loading. The biochemical model indicates a possible mechanism linking ATP levels to codon optimality effects that is consistent with observations from GTEx and mouse data. We then set out to test through perturbation analyses whether this link was causal or merely correlative. The GTEx dataset, consisting of post-mortem samples, provides data for which Nature cruelly performed such a perturbation experiment. At death, respiration and blood circulation cease, which deprives the body’s cells of oxygen and impairs mitochondrial ATP synthesis. Moreover, the GTEx samples were stabilized at different times after death, or ischemic times, allowing us to study the effects of varying degrees of oxygen deprivation. We found the COMD coefficient to increase with ischemic time adjusting for age and tissue (multivariate analysis Fig. 3A, stratification Fig. 3B, Fig. S6-10). These results, which show that the effect of codon optimality on mRNA stability is amplified upon oxygen deprivation, further support our model.

**Fig. 3:**
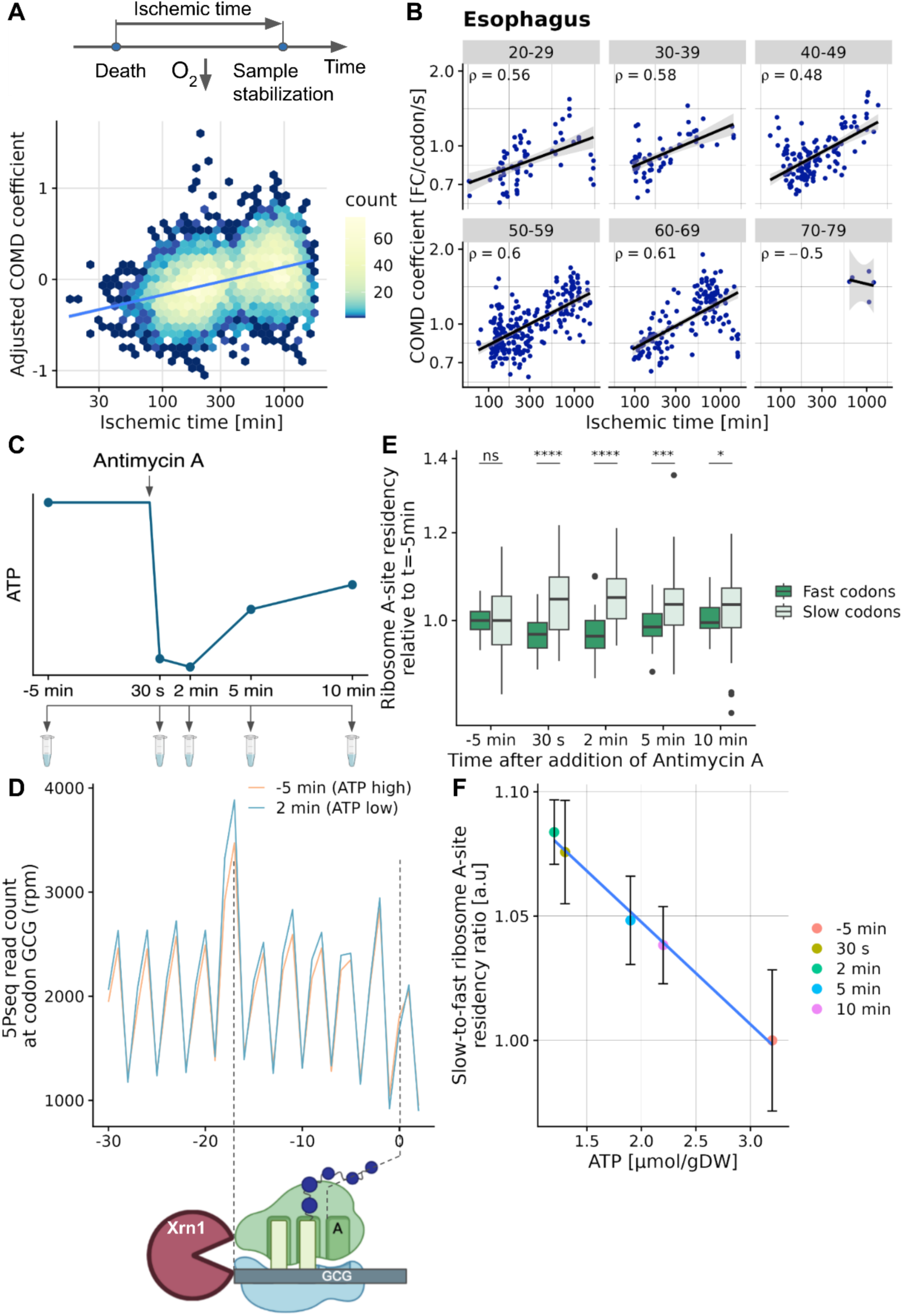
Differences in decoding of fast and slow codons depend on intracellular ATP concentration. **A** COMD coefficient adjusted for age and tissue (Methods) against ischemic time (Spearman’s rho = 0.41 *P* < 10^−15^). Blue line marks linear trend obtained by linear regression. **B** COMD coefficient against ischemic time for esophagus samples grouped by age category (Spearman’s rank correlation rho is statistically significant (*P*<0.05) for all age categories but 70-79). **C** Sampling design of 5P-seq profiling time course following addition of the cellular respiration inhibitor Antimycin A on yeast cells. **D** Number of 5P-seq reads (in reads per million) per position relative to the GCG codons for the time points with maximum (−5 min) and minimum (2 min) intracellular ATP concentration. 5PSeq maps the 5’ends of mRNAs co-translationally degraded by Xrn1, which are located 17 nucleotides 5’ of the ribosome A site [25]. Hence, the peak located 17 nt 5’ of GCG codons is consistent with GCG being a non-optimal codon in yeast. This peak is amplified at low ATP concentration (2-minute time point) in agreement with the biochemical model (Fig. 2). **E** Distribution of ribosome A-site residencies inferred from 5PSeq for slow and fast codons (defined using first and fourth quartiles of occupancy in unperturbed cells). Values are expressed in fold-change relative to median occupancy in unperturbed cells (Methods). P-values were obtained from double-sided Wilcoxon rank sum tests (* < 0.05, ** < 0.01, *** < 0.001, **** < 0.0001). **F** Difference in ribosome A-site residency for each time point between slow and fast codons and its corresponding reported intracellular ATP concentration [26].

However, ischemic times in the GTEx dataset span hundreds of minutes. It cannot be excluded that mechanisms other than intracellular ATP deprivation [27], including tRNA regulatory response, could be occurring during these times. To address this concern, we next performed an experiment to assay the short-term response of ribosome dwell time to ATP deprivation. To this end, we profiled ribosome occupancies of *S. cerevisiae* cultures following exposure to antimycin A, a cellular respiration inhibitor (Fig. 3C). Ribosome occupancies profiling was performed using 5’P sequencing (5PSeq) [28,29]. Importantly, 5PSeq, which does not require in vitro incubation with RNase, allowed us to obtain measurements in the first few minutes after drug application. We estimated the relative ribosome residency on A-site codons by adjusting 5PSeq coverage for the 17-nt shift due to ribosome protection as well as controlling for gene coverage, distance to the start codon, and sequencing depth (Fig. 3D for raw data around a non-optimal codon, Methods). As expected, these estimates of relative ribosome residency on A-site codons were consistent with previously reported decoding times and codon optimality in yeast (Fig. S11). Notably, A-site residency ratios between optimal and non-optimal codons negatively correlated with intracellular ATP concentration over the time course (Fig. 3E, F).

These results are consistent with ATP concentration modulating the effect of codon optimality on translation and translation-related mRNA degradation. Furthermore, these observations made within a few minutes upon ATP deprivation, cannot be explained by a transcriptional response of tRNA abundance.

Our findings reveal a fundamental link between metabolism, gene expression, and gene sequence, the consequences of which need to be explored. As a first step in this direction, we asked whether this phenomenon could constrain tissue-specific mRNA isoform regulation. To this end, we considered cassette exons, i.e. exons located in between two other exons, exhibiting a tissue-specific inclusion pattern. In sun-exposed skin, a tissue with a high COMD coefficient and low mitochondrial activity, we found that cassette exons that were more often included used more optimal codons compared to cassette exons that were more often skipped (Fig. 4A). The opposite was observed in the cerebellar hemisphere, a tissue with a low COMD coefficient and high mitochondrial activity. Furthermore, the trend generalized to tissues with intermediate COMD coefficient values (Fig. 4B). Hence, these observations reveal an unanticipated codon usage bias for tissue-specific exons. Moreover, our model provides a mechanistic explanation for this: non-optimal codons are relatively less likely to trigger mRNA degradation in tissues with high cellular energy. Therefore, splice isoforms using non-optimal codons are comparatively more stable in tissues with high cellular energy, which is reflected in an increased abundance of cassette exons using slow codons in the mRNA pool. Perhaps, even, these exons have evolved to use codons suited to the metabolic state of the tissue they are expressed in.

**Fig. 4:**
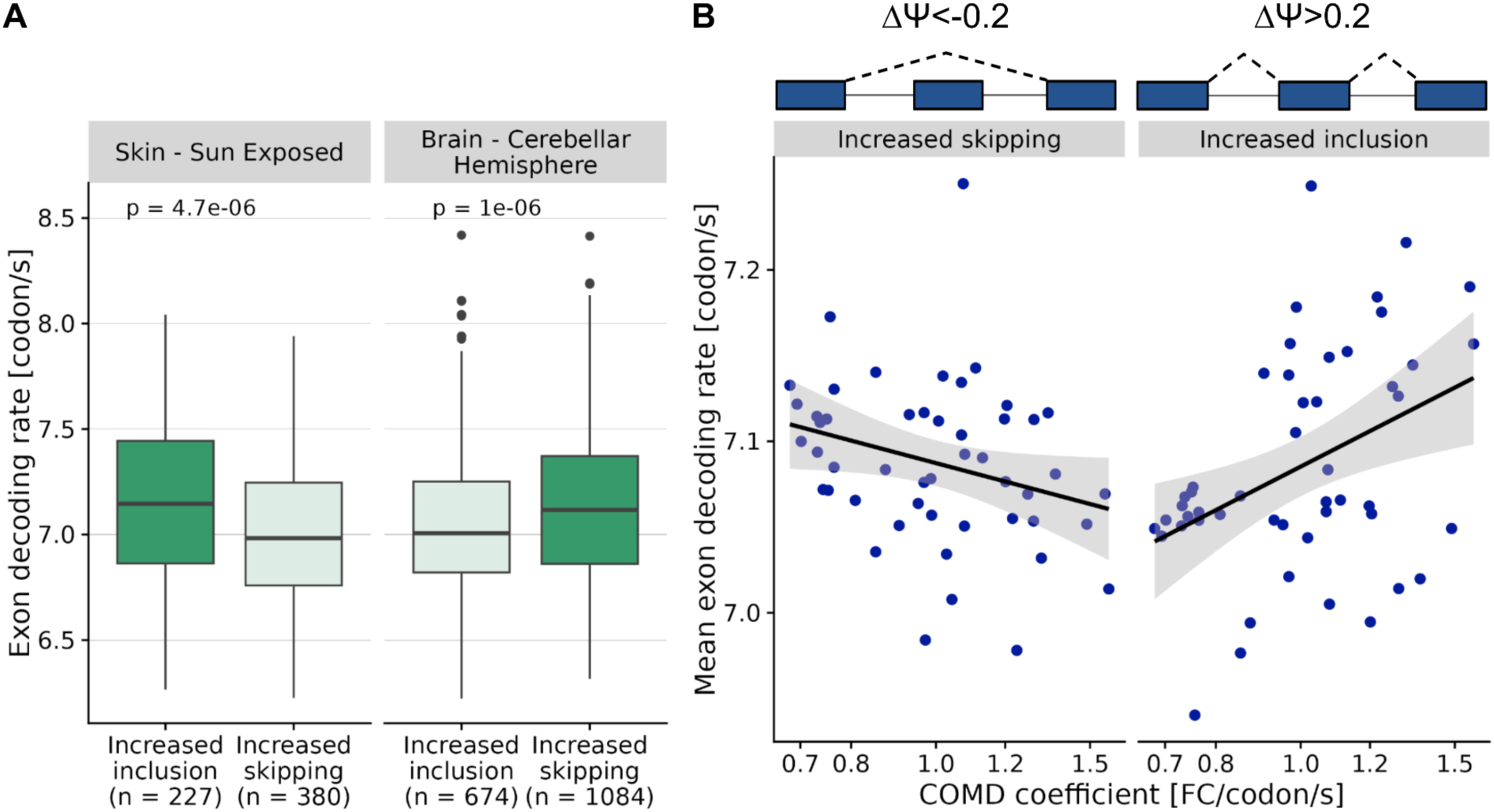
Codon optimality of cassette exons increasingly included or skipped in a tissue relates to its COMD coefficient. **A** Distribution of the decoding rate of cassette exons that are increasingly included (ΔΨ > 0.2) or skipped (ΔΨ < −0.2) in sun-exposed skin (left) and cerebellar hemisphere (right) compared to other tissues. Exon decoding rates are expressed in codons per second and computed using codon decoding rates measured in HEK293 cells (Methods). *P*-values were obtained from a Wilcoxon rank sum test. **B** Codon decoding rate average of increasingly skipped (left) and included (right) exons per tissue against the tissue’s COMD coefficient. Spearman’s rank correlation rho is statistically significant for both panels (*P* = 0.018 and *P* = 0.007, one-sided *P*-value).

## Discussion

Taken together, our results show that cellular energy regulates the effect of codon optimality on mRNA stability and translation. Codon optimality matters more in conditions of scarcer energy, such as tissues with low mitochondrial activity, older age, oxygen deprivation, or exposure to specific drugs. This is explained by a nonuniform effect of the abundance of energy-carrier molecules on codon decoding, which occurs independently of tRNA abundance regulation. Our results provide furthermore a mechanistic basis for previous observations of attenuated effects of codon optimality in brain and testis of the fruit fly [6,9], and, as ATP concentration is maximal at the G2/M phase [30], on attenuated effects of codon usage in proliferative cells in absence of tRNA regulation [7].

## Methods

### GTEx Exonic and intronic read counts

The BAM files for 7,778 RNA-Seq samples, the gene-level and the transcript-level TPM (transcript per million) values, as well as the sample annotation of the GTEx v6 dataset, genome build hg19, were downloaded from the GTEx portal on June 12, 2017, under accession number dbGaP: phs00424.v6.p1. This RNA-Seq dataset is paired-end and unstranded. Exonic coordinates of all protein-coding genes located in standard chromosomes were extracted from the GENCODE annotation (Frankish et. al. 2021), release 19. Exonic and intronic read counts were obtained as recommended by Gaidatzis et al. [14]. Specifically, exonic coordinates were flanked on both sides by 10 nt and were grouped by gene. Intronic coordinates were obtained by subtracting the exonic coordinates from the gene-wise coordinates. For each gene, exonic and intronic read counts were obtained using the summarizeOverlaps function from the GenomicAlignments package (Lawrence et. al. 2013) v. 1.28.0 with the mode parameter set to “IntersectionStrict” and the inter.feature parameter set to FALSE to consider only reads that fully fall within the desired genomic regions. Moreover, to be robust against noisy estimates based on low read counts, in each sample genes with TPM < 1 were ignored (read counts set as missing values). Finally, for each gene and each sample, the log-transformed exonic-to-intronic read count ratio *y* was computed using pseudocounts of 1:

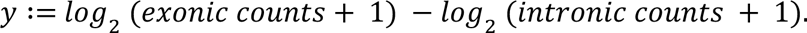

### Relative mRNA half-life

Next, we attributed those *y* values to the major isoform for each sample, whereby the major transcript isoform was taken as the transcript isoform with highest median TPM value across samples of the same tissue. Transcripts with missing values in more than one third of samples were discarded. For the remaining transcripts, tissue-specific log-transformed exonic-to-intronic read count ratio were calculated by taking the median of the *y* values over all samples from the same tissue. We further filtered out transcripts with missing values in more than 15% of the tissues.

Exonic and intronic read counts depend on gene-specific biases including exonic and intronic length and GC-content. Following Gaidatzis et al. [14], these biases are mostly multiplicative and therefore cancel out when considering ratios between samples. We therefore use *y* differences as estimates for mRNA half-life log-ratios between pairs of samples or tissues. Following the same logic, the log2 relative mRNA half-life 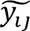 for transcript *i* in sample or tissue *j* was defined relatively to the average across tissues or samples, respectively, i.e.:

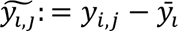

The entire procedure was applied at two levels of tissue annotation granularity: the GTEx major tissues on the one hand, and the GTEx subtissues on the other hand.

### Transcript sequence features

Transcript sequences were retrieved using pyranges v0.0.84, kipoiseq v0.4.1 from the release 19 of the GENCODE annotation of the human genomic sequence GRCh37/hg19. Only coding sequences starting with the start codon AUG were considered.

### Codon effects

The estimated codon effect, *sk,j*, was obtained for each codon *k* and tissue *j* separately by fitting a univariate linear regression of the type:

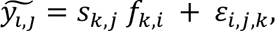

where *f_k,i_*, is the proportion of codon *k* in the coding sequence of transcript *i*. To this end, we used scikit-learn v0.22.2.

### COMD coefficient

The COMD coefficient was obtained for each tissue or sample *j* by fitting a linear regression of the type:

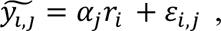

where *αj* is the COMD coefficient for the sample or the tissue *j* and *ri* is the geometric mean of the codon decoding rates of transcript *i* in HEK293.

### Generation of spliced and unspliced read counts ratio for the mouse dataset

The BAM files and valid barcodes for the 28 scRNA-Seq samples (using droplet-based 10x Genomics Chromium protocol) of 3-month old mice of the Tabula Muris Senis atlas dataset [19] were downloaded from the public AWS S3 bucket (https://registry.opendata.aws/tabula-muris-senis/). The reference genome used to generate the BAM files did not contain mitochondrial-encoded genes, therefore they were not considered for our analysis. Sample annotations were downloaded from the Gene Expression Omnibus under accession code GSM4505404, specifically the file GSM4505404_tabula-muris-senis-droplet-official-raw-obj.h5ad.

Loom files for each sample containing raw spliced and unspliced counts were obtained by running the velocyto command-line tool ([31], v0.17.17). In contrast to bulk RNA-Seq, UMI-based scRNA-Seq (unique molecular identifier) allows identifying whether reads originated from the same transcript. The velocyto tool makes use of this and collectively marks reads of the same transcript as unspliced if one of them aligns to an intronic region or exon-intron junction. Conversely, if all reads of the same UMI solely align to exonic regions, they are marked as spliced reads. Equivalently to exonic and intronic reads, unspliced reads are a proxy for premature mRNA and spliced reads a proxy for mature mRNA, therefore their ratio is a proxy for mRNA stability [14].

In order to have enough cells for pseudo-bulking, we filtered out cell types (“cell_ontology_class”) that had less than 150 cells using scanpy [32], v1.8.2). For the remaining 45 cell types, we computed pseudo-bulk aggregates by summing all counts of cells per cell type for spliced and unspliced counts independently. To only consider genes that are expressed across multiple cell types, we filtered out genes with less than 3,000 counts shared across spliced and unspliced counts and all cell types. We normalized both spliced and unspliced counts by dividing them by the total number of spliced and unspliced counts, respectively, over all genes per cell type. Per gene spliced-to-unspliced ratios were computed as log10(spliced counts + 1) - log10(unspliced counts + 1). Spliced-to-unspliced ratios were centered across cell types for each gene. As gene expression, we used the normalized and log-transformed spliced counts.

### Transcript sequence features for the mouse dataset

Transcript features for mouse were retrieved as described for human. For mouse, we used release 25 of the GENCODE annotation [33] of the mouse genomic sequence GRCm38/hg38.

### Major transcript isoform selection for the mouse dataset

Since the 10x single cell technology does not produce full length transcript coverage, the major transcript of a gene was defined as the transcript with the highest support. We considered the tags of each transcript (GENCODE, release 25) and calculated the support by summing up the following categories: being part of the GENCODE basic annotation (tag “basic”), being tagged as the principal isoform according to APPRIS database (tag “appris_principal_1”, [34]), being a member of the consensus CDS gene set (tag “CCDS”, [35]), and having an Ensembl transcript support level of 1 (“transcript_support_level”). A transcript could therefore have a support between 0 and 4 and for each gene we chose the transcript with the highest support. If there were ties, one transcript was randomly chosen.

### Gene set enrichment analysis

Gene set enrichment analysis [18] was performed on the genes scored by their Spearman correlation between their TPM values and the COMD coefficient across tissues using GSEApreranked with package gseapy v1.0.0. Only genes with TPM > 1 in all tissues were considered. Within-tissue correlation between the COMD coefficient and gene expression (TPM) followed by GSEA was computed as before. Only genes with TPM > 1 in at least two thirds of the samples were considered.

In mouse, we considered genes whose log-transformed and normalized spliced counts were greater than 0 across all cell types (6,051 genes). As in human, gene set enrichment analysis was performed on the genes scored by their Spearman correlation.

For all GSEA analyses, only pathways with FDR ≤ 0.01 were considered significantly enriched.

### Individual’s COMD coefficient

We fitted a linear regression predicting the COMD coefficient of each sample as the sum of an individual’s effect (which we defined as the individual’s COMD coefficient) and a tissue effect.

### Overview of the elongation cycle model

We developed a simplified steady-state model of one translation elongation cycle to assess how variations in ATP and GTP abundance can change the rate of ternary complex loading into the ribosome for given tRNA and ribosome concentrations. We modeled the 2-1-2 pathway of E-site tRNA release, because, unlike the 2-3-2 pathway, it includes the state of the ribosome in which both the A-site and the E-site are free [36], which is the substrate for the Ccr4-Not complex [3].

In our simplified model, we considered the ribosome in two states: free A-site or occupied A-site. The free A-site state corresponds to the point where the ribosome is ready to accept its cognate ternary complex. In this state the ribosome has finished translocation, the P-site is occupied and both the A-site and the E-site are free.

The occupied A-site state represents the point where the ribosome is ready to translocate. In this state the tRNA has already been accommodated into the A-site (following GTP hydrolysis) and the new peptide bond formed. In this state both the A-site and the P-site are occupied and the E-site is free.

The transition between the free and the occupied A-site states described above is characterized by a series of intermediate states, such as peptide-bond formation and ribosome conformation changes (reviewed by Dever et al. [22]), that do not depend on the variables of interest: tRNA, ribosome, GTP and ATP concentrations. Therefore, under changes in these variables, the rate of transition between such intermediate states gives a constant contribution to the rate of transition between the free and occupied A-site states. Following from this, we considered the following reactions: tRNA aminoacylation, ternary complex formation, ternary complex loading, ribosome translocation (Fig. 2E).

### tRNA aminoacylation

tRNAs are charged with amino acids by the aminoacyl-tRNA-synthetase in a two-step reaction, where ATP is hydrolyzed to AMP and one amino acid is loaded into the tRNA (reviewed by Gomez and Ibba [23]). We modeled tRNA aminoacylation with an overall irreversible reaction:

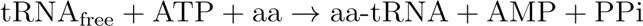

where tRNA_free_ represents the pool of uncharged tRNAs available to be aminoacylated. Assuming law of mass action, we modeled the rate of tRNA aminoacylation as:

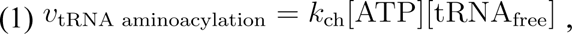

where *k*_ch_ is the rate constant of the tRNA aminoacylation reaction under a given concentration of amino acids.

### Ternary complex loading

The ternary complex, composed of one aminoacyl-tRNA, one GTP, and one eukaryotic translation elongation factor 1A (eEF1A) binds in the A-site of the ribosome, where the aminoacyl tRNA is accommodated after the hydrolysis of GTP followed by the release of eEF1A-GDP [22]. We modeled the TC binding to the A-site of the ribosome and the subsequent tRNA accommodation as a single irreversible reaction which we termed TC loading:

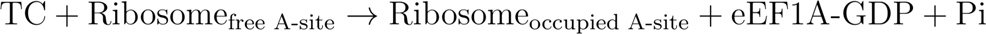

We assumed law of mass action kinetics, i.e:

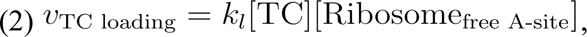

where *k*_l_ is the rate constant of the TC loading reaction.

### Ribosome translocation

Translocation of the ribosome requires the binding of eEF2-GTP to the A-site of the ribosome which is followed by the hydrolysis of GTP and subsequent release of eEF2-GDP. After translocation, the deacylated tRNA on the E-site is released from the ribosome [22]. We modeled this process as a single irreversible reaction combining the recharging of eEF2 with GTP and its subsequent binding to the ribosome A-site followed by GTP hydrolysis, which results in ribosome translocation. Furthermore, we combined together ribosome translocation and the release of tRNA from the E-site:

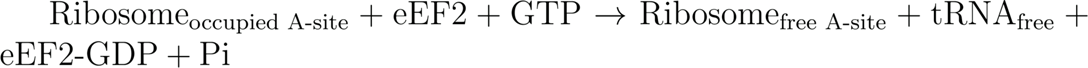

We assumed law of mass action kinetics, i.e.:

(3) *v*_translocation_ = *k*_tr_ [GTP][Ribosome_occupied A-site_], where *k*_tr_ is the rate constant of the translocation reaction under a constant concentration of eEF2.

### Ternary complex formation

Aminoacyl-tRNAs (aa-tRNAs) are bound to the ribosome in a ternary complex (TC) with GTP and eEF1A (elongation factor 1A). eEF1A is charged with GTP by the exchange factor eEF1B [22]. We modeled the eEF1A charging with GTP and subsequent binding to aa-tRNA into a single reversible reaction:

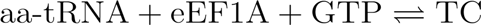

We assumed law of mass action kinetics, i.e.:

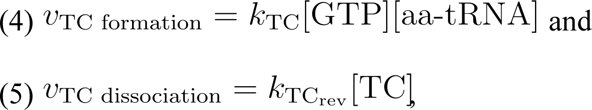

where *k*_TC_ and *k*_TCrev_ correspond to the rate constant of the TC formation and TC dissociation reactions respectively under a constant concentration of eEF1A.

### Steady-state solution

Assuming steady state, we symbolically solved the resulting system of equations in Wolfram Mathematica v13.0.0.0. This led to the following expression for the rate of TC loading as a function of ATP, GTP, tRNAtotal, Rtotal and all rate constants considered above:

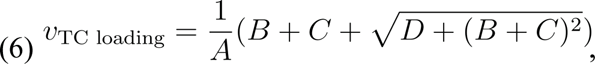

where:

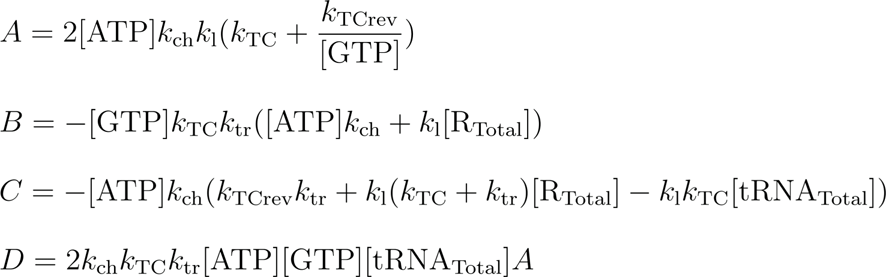

the total number of ribosomes is:

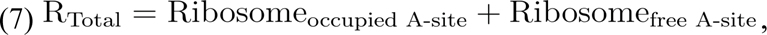

and the total number of tRNAs is:

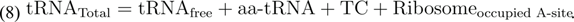

The order of magnitude of each rate constant was estimated based on previous reports and mostly taken from *S. cerevisiae* when possible. To ensure our results are not qualitatively sensitive to the choice of rate constant, we considered multiple values spanning 2-3 orders of magnitude around the ones based on literature.

In *S. cerevisiae*, the rate of tRNA aminoacylation is in the order of magnitude of 1 tRNA per second [37] and ATP concentration can vary between 2 and 6 mM [38]. Following Equation 1, we can infer that kch ≈ 10^3^ s^−1^M^−1^.

From [39] the rate constant of ternary complex loading into the ribosome in *S. cerevisiae* is kl ≈ 10^7^ s^−1^M^−1^.

The rate of translocation for one ribosome is vtranslocation ≈ 10^2^ s^−1^ [39]. Given the concentration of GTP ≈ 0.1 mM [40], BNID 101420 [41] and considering the previously derived expression for translocation rate velocity, kt ≈ 10^6^ s^−1^M^−1^.

The rate of ternary complex association for one aminoacyl-tRNA was estimated to be vTC ≈ 10^1^s^−1^in *E. coli* [42] assuming a constant concentration of the required elongation factor (EF-Tu) in its normal range. Given the concentration of GTP ≈ 0.1mM and the previously derived expression for the rate of TC formation, vTC, kTC ≈ 10^5^ s^−1^M^−1^.

The concentration of tRNAs in yeast commonly varies between 0.1 μM and 1 μM [39]. The total concentration of ribosomes is in the range 1-10μM [39]. Assuming that between 1% and 10% of the ribosomes translate one specific codon, the concentration of ribosomes translating it is in the range 0.01-1 μM. On figures 2F,G the slow codon has a ratio between the total number of tRNAs and total number of ribosomes of 1:4, and their concentration is 0.2μM and 0.8μM respectively. For the fast codon, the ratio between the total number of tRNAs and total number of ribosomes is 4:1, and their concentration is 0.4 μM and 0.1 μM respectively.

GTP concentration is found to be one order of magnitude below ATP concentration according to data from [40] and [38]). As an approximation, we assumed a GTP concentration proportional to ATP concentration in 1:10 ratio.

Finally, the relative difference between the minimal and maximal ATP concentrations of the fast-to-slow TC loading rate ratio was computed using Equation 6, for combinations of rate constants between 10^2^−10^4^ for kch, 10^7^−10^8^ for kl, 10^4^−10^6^ for kTC, 10^5^−10^7^ for ktr, and 10^−3^−10^−1^ for kTC rev. For each interval, we considered all possible values in steps of 0.1 in the order of magnitude. The fast-to-slow TC loading rate ratio was then computed for every parameter value combination (over 2.1 million combinations).

### Ischemic time analysis

The sample ischemic time in minutes is defined by GTEx as “the interval between actual death, presumed death, or cross clamp application and final tissue stabilization” and was obtained from the GTEx sample annotation file, column “SMTSISCH”. The ischemic time was compared to the COMD coefficient adjusted for age and tissue, defined as the residual of the linear regression predicting the sample COMD coefficient as a linear combination of the age and a tissue effect.

### Yeast strains and culture conditions

The strain used in this study was BY4741 Mat a ura3 met1 his3 leu2 background (referred to as wild type). *S. cerevisiae* were grown in trehalose-containing medium as the non-fermentable carbon source at 30°C. Trehalose-containing medium includes 20 g/l trehalose, 6.7 g/l YNB + (NH4)2SO4 (yeast nitrogen base without amino acids; Difco), amino-acid supplements at a final concentration of 100 mg/l to complement the auxotrophies of the strains. The medium was buffered at pH 4.8 by adding 14.6 g/l succinic acid and 6 g/l NaOH [26]. Pre-cultures were grown overnight in 250 mL flasks and agitated at 150 rpm. The next day, pre-cultures were diluted to OD600 = 0.05 and grown until an OD600 of ∼0.5 was reached. To block mitochondrial function, antimycin A was added to a final 2 μg/ml to the medium and incubated for 0, 2,5, and 10 min [26]. Cells were fast spun down for 15 seconds at 13,000 g in a microcentrifuge and flash-frozen in liquid nitrogen.

### HT-5PSeq library preparation

HT-5PSeq libraries were prepared as previously performed (Zhang & Pelechano, 2021). Briefly, 6 μg RNA was used for DNase treatment, then DNA-free total RNA were directly ligated with RNA/RNA oligo containing UMI (RNA rP5_RND oligo). Ligated RNA was reverse transcribed and primed with Illumina PE2 compatible oligos containing random hexamers and oligo-dT. RNA in RNA/DNA hybrid was depleted by sodium hydroxide within 20 min at 65°C incubation. Ribosomal RNAs were depleted using DSN (Duplex-specific nuclease) with the mixture of ribosomal DNA probes. Samples were amplified by PCR and sequenced in an Illumina NextSeq 2000 instrument using 55 sequencing cycles for read 1 and 55 cycles for read 2.

### HT-5PSeq analysis

HT-5PSeq reads were trimmed 3’-sequencing adapter using cutadapt V1.16 [43]. The 8-nt random barcodes on the 5’ ends of reads were extracted and added to the header of the fastq file as the UMI using UMI-tools. 5’P reads were mapped to the *S. cerevisiae* reference genome (SGD R64-1-1) using STAR 2.7.9a with the parameter --alignEndsType Extend5pOfRead1 to exclude soft-clipped bases on the 5’ end. After removing PCR duplicates by UMI-tools 1.0.0 [44], analysis of 5’ ends positions was performed using the Fivepseq package [45], including the relative distance to start and stop codons. Specifically, the unique 5’mRNA reads in biological samples were merged.

### 5PSeq modelling

We considered all 5PSeq reads located 17 nt 5’ of in-frame codons. To avoid possible confounding effects due to translation initiation and termination, we did not consider the start codon and the second codon, nor we considered the stop codon and its preceding codon. For robustness, we further did not consider genes with less than 10 reads mapping to all considered codons. Furthermore, genes encoded in mitochondrial DNA were discarded because they use a different genetic code.

We started by isolating the contribution of the codon-specific ribosome dwelling to the 5PSeq coverage from factors independent of translation elongation. The 5PSeq read coverage can depend on how frequently the corresponding mRNA is degraded, its expression and how frequently translation is initiated. Furthermore, as Xrn1 follows the last translating ribosome, the position of the codon in the coding sequence could potentially impact the 5PSeq coverage. Hence, we modeled the reads in gene *g* and distance *d* from the start codon in order to isolate the effect of the A-site codon on the 5PSeq reads from gene, position and sequencing depth effects. For each sample (belonging to a batch and time point) we modeled the read coverage *y_g,d_* on gene *g* at distance *d* from the start codon (adjusting for the 17 nt shift) using a generalized linear model with a negative binomial distribution and the log link function:

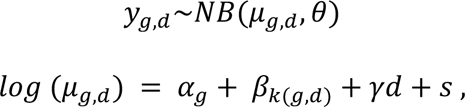

where α*_g_* is a gene effect, β*_k(g,d)_* is the relative ribosome residency for the A-site codon located at distance *d* on gene *g*, and *s* (or size factor) is a sample-specific intercept.

The model was fitted using the package *statsmodels* v0.12.0. Due to memory limitations, the model was fitted separately on three thirds of the data for each sample, and the coefficient estimates averaged over the three thirds.

To define fast and slow codons for the yeast 5PSseq time course, we considered the samples harvested prior to application of the drug. Next we averaged the relative ribosome residencies β_*k*_ for every codon *k* across these samples and defined the fast codons as the codons from the first quartile and the slow codons as the codons from the fourth quartile.

To relate intracellular ATP concentration with the 5P-seq readouts at time point 0 while accounting for centrifugation and sample stabilization of the 5PSeq protocol, we considered the intracellular ATP concentrations reported by [26] at 30 seconds. For the later time points (2 min, 5 min, 10 min), we used the very same time points of Walther.

### Cassette exons

Exon annotations and percent-spliced-ins (PSI) were obtained from the ASCOT annotation [46]. We filtered for cassette exons according to ASCOT (i.e. feature cassette_exon=Yes) that belong to genes of the consensus CDS gene set of the GENCODE release 25 (build GRCh38). We further restricted the analysis to such cassette exons that fully overlap a coding sequence. This resulted in 29,417 cassette exons. The decoding rate of each coding cassette exon was then calculated as the geometric mean of the HEK293 decoding rates [17] of the codons fully contained in the exon. A cassette exon was considered tissue-specifically differentially spliced if its PSI was at least 20 percent points above or below its average PSI across tissues.

## Ethics approval and consent to participate

Not applicable.

## Consent for publication

Not applicable.

## Availability of data and materials

The 5PSeq data has been deposited in NCBI’s Gene Expression Omnibus [47] and will be accessible through GEO Series accession number GSE216524 upon acceptance of the manuscript. Authorized researchers can access the BAM files for 7,778 RNA-Seq samples, the gene-level and the transcript-level TPM (transcript per million) values, as well as the sample annotation of the GTEx v6 dataset from the GTEx portal. The BAM files of the Tabula Muris Senis atlas dataset [19] can be downloaded from the public AWS S3 bucket (https://registry.opendata.aws/tabula-muris-senis/). The code developed for the analysis is available at https://github.com/gagneurlab/Cellular_energy_codon_analysis.

## Authors’ contributions

Conceptualization: P.T.S, V.P., J.G.; Methodology: P.T.S., Y.Z., E.T., L.D.M., V.P., J.G.; Software: P.T.S, Y.Z., E.T., L.D.M.; Formal analysis: P.T.S., E.T., Y.Z; Investigation: P.T.S., Y.Z.; Resources: Y.Z., V.P.; Data Curation: P.T.S., Y.Z., E.T., L.D.M.; Writing – original draft: P.T.S., J.G..; Writing - review & editing: P.T.S., Y.Z., V.A.Y., V.P., J.G.; Visualization: P.T.S., Y.J., V.A.Y., J.G..; Supervision: V.P., J.G.; Project administration: V.A.Y., J.G.; Funding acquisition: V.P., J.G.

## Acknowledgements

We would like to thank Diego Núñez, Thomas Becker and Roland Beckmann for advice and feedback on the manuscript. Furthermore, we are grateful to Alexander Karollus, Leonhard Wachutka, Felix Brechtmann and all the remaining members of the Gagneur Lab for the discussions.

