## Supplementary Material for "Cellular energy regulates mRNA translation and degradation in a codon-specific manner"

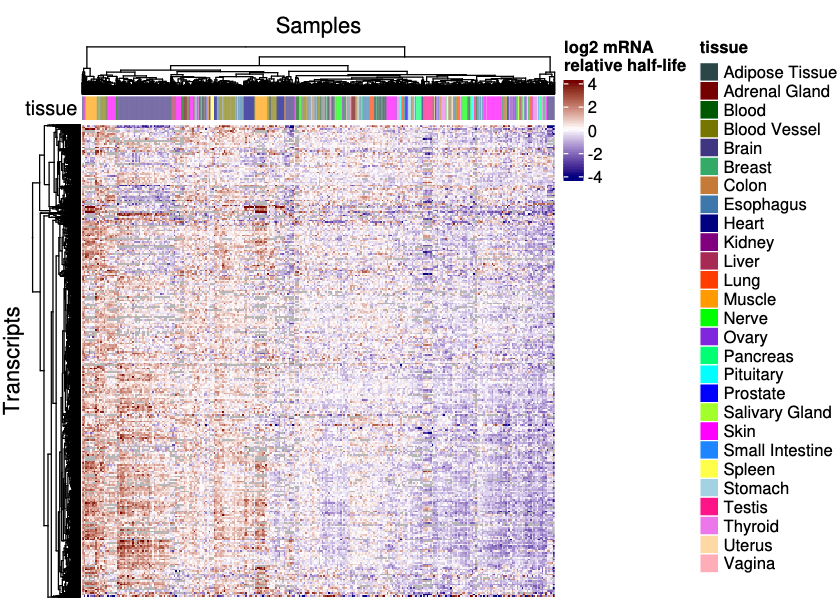

**Figure S1:** **Relative mRNA half-lives across samples and tissues.** The columns represent transcript major isoforms expressed in at least ⅔ of the samples. log_2_ relative mRNA half-life was saturated at -4 and 4. Each sample is colored by the major tissue it belongs to.

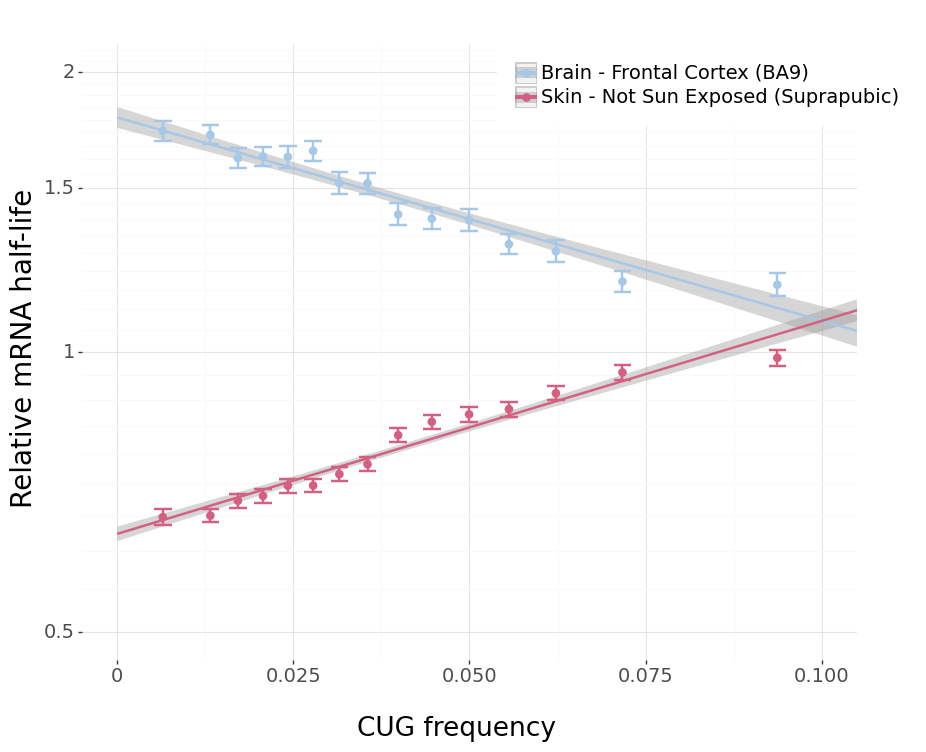

**Figure S2: Frequency of the optimal codon CUG in the coding sequence relates to changes in mRNA half-life differently depending on the tissue.** The mean and standard error bars of relative mRNA half-life are plotted for each group of mRNAs within equally-sized bins of CUG frequency. Transcripts with low usage of CUG exhibit large half-life fold changes between Brain - Frontal Cortex (BA9) and Skin Not Sun-Exposed. In contrast, transcripts with high usage of CUG exhibit a mild half-life fold change. The result of a linear regression of relative mRNA half-life on the frequency of CUG is plotted for each tissue. The slope of each line corresponds to the estimated codon effect of CUG for these tissues reported in the heatmap of Figure 1B.

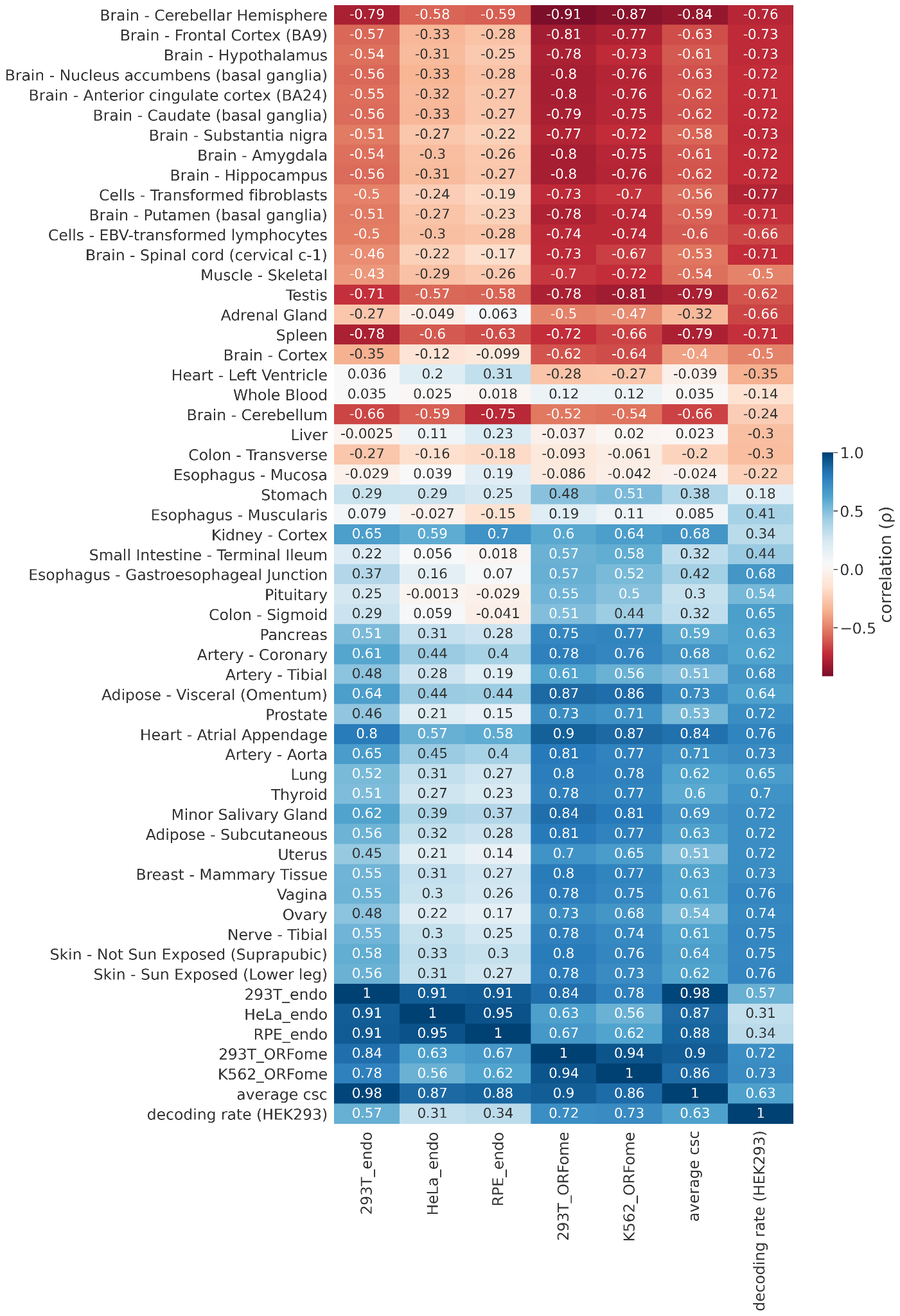

**Figure S3: Codons change in their associations with relative mRNA half-life according to their optimality and decoding rate.** Each cell of the heatmap represents the correlation between the estimated codon effects for each tissue (rows) and different metrics of codon optimality and decoding rate computed in different cell lines (columns). The labels 293T_endo, Hela_endo, RPE_endo, 293T_ORFome, K562_ORFome correspond to the CSC (codon stability coefficient) in the different cell lines (Hela, 293T, RPE, K562) computed either from endogenous mRNAs (endo) or constructs (ORFome) and were obtained from [48]. The decoding rate in HEK293 was obtained from [17].

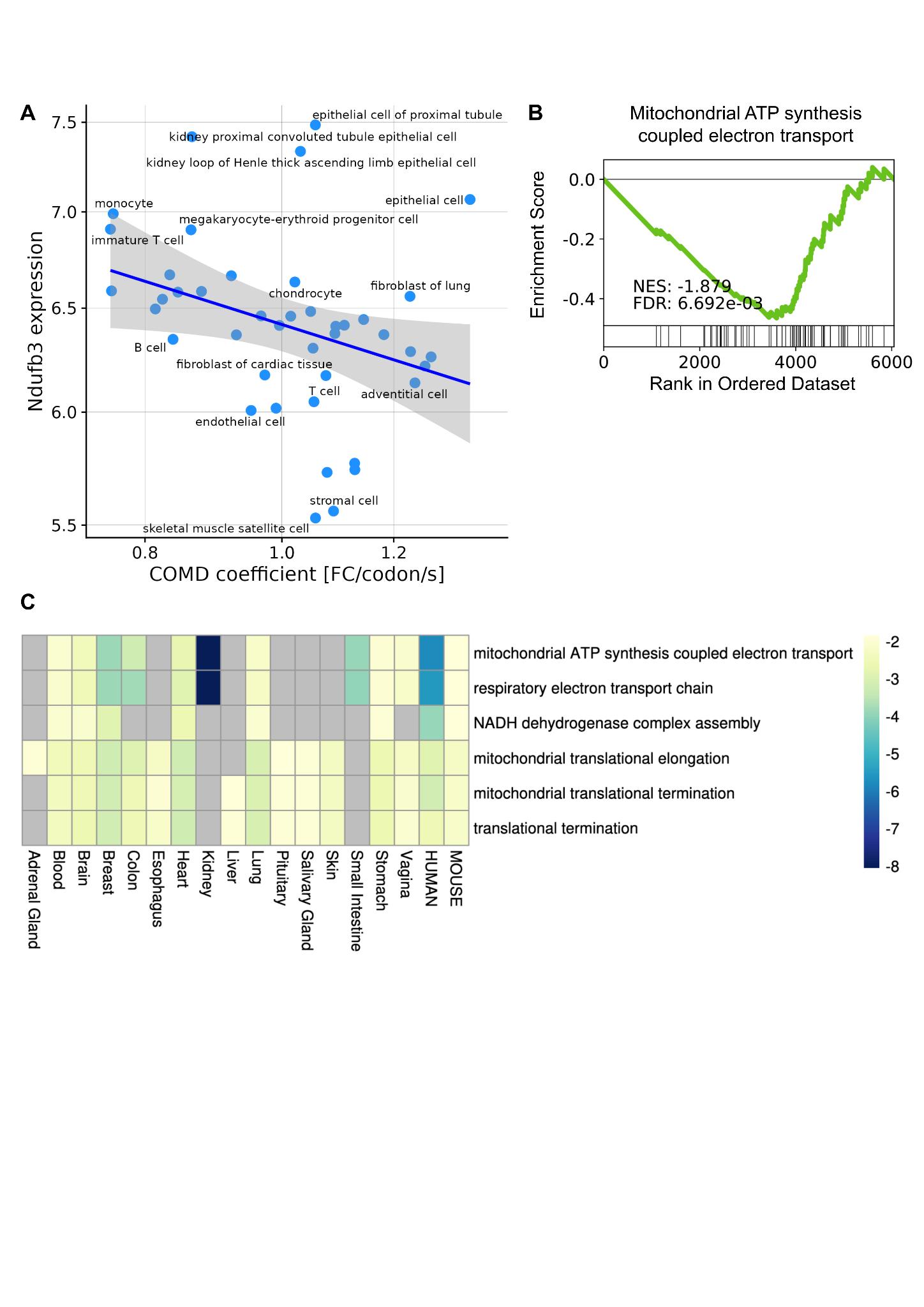

**Figure S4: Mitochondrial-related pathways are enriched for differences in COMD coefficient across and within tissues in human and across cell types in mouse. A** Relationship between the COMD coefficient and the expression of *Ndufb3*, a gene that encodes a subunit of mitochondrial respiratory complex I, across cell types in mouse (Spearman’s rho = -0.44, *P* = 0.0034). **B** Gene set enrichment analysis for the biological process “mitochondrial ATP synthesis coupled electron transport” in mouse. **C** GSEA normalized enrichment score for significant pathways of genes correlated with variations in the COMD coefficient across samples from the same tissue, between tissues in human and between tissues in mouse. Pathways shown are restricted to the ones commonly found in human and mouse.

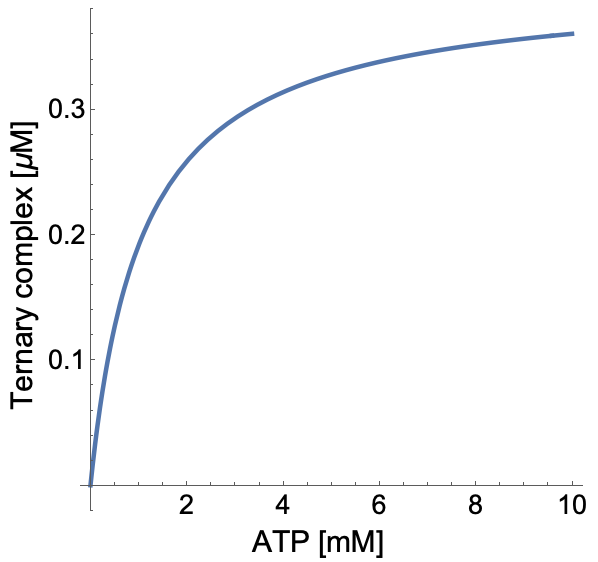

**Figure S5: Ternary complex concentration increases asymptotically with ATP concentration.** Ternary complex against ATP concentration for the same rate constant values as in Figure 2F,G and tRNA and ribosome concentrations of 0.4μM and 0.1μM respectively.

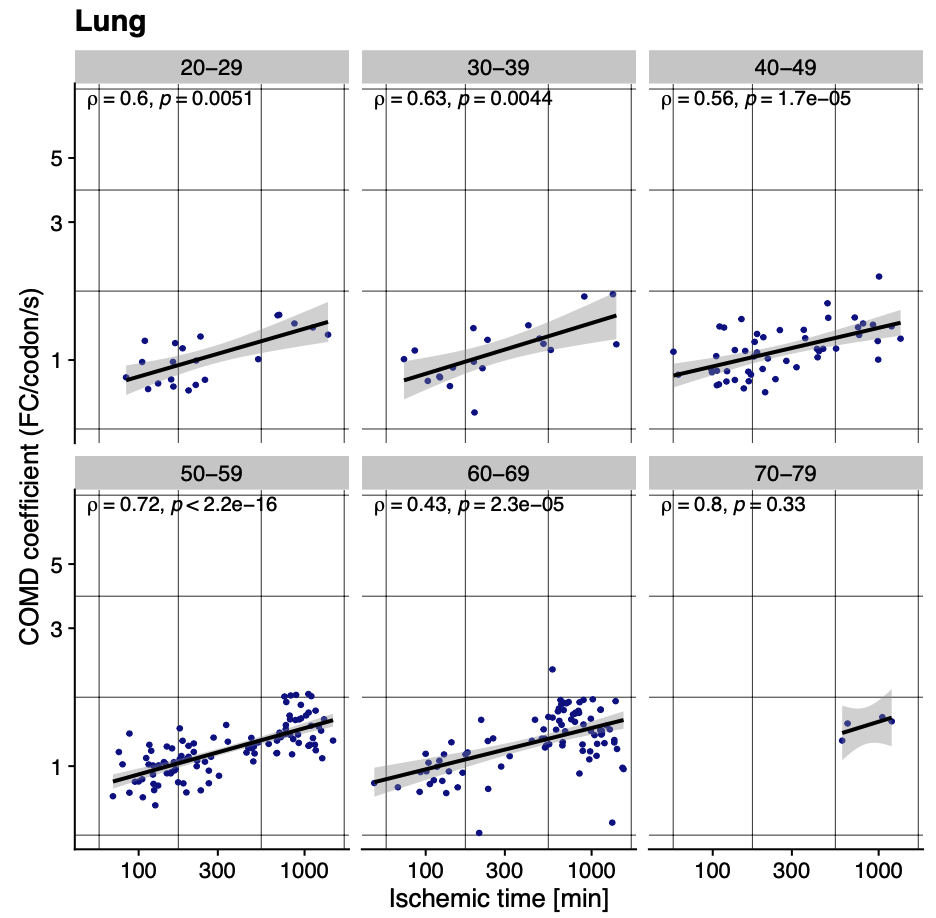
**Figure S6: The COMD coefficient associates with ischemic time for individuals of similar age in Lung.**  COMD coefficient against ischemic time (min) for Lung in different age groups.

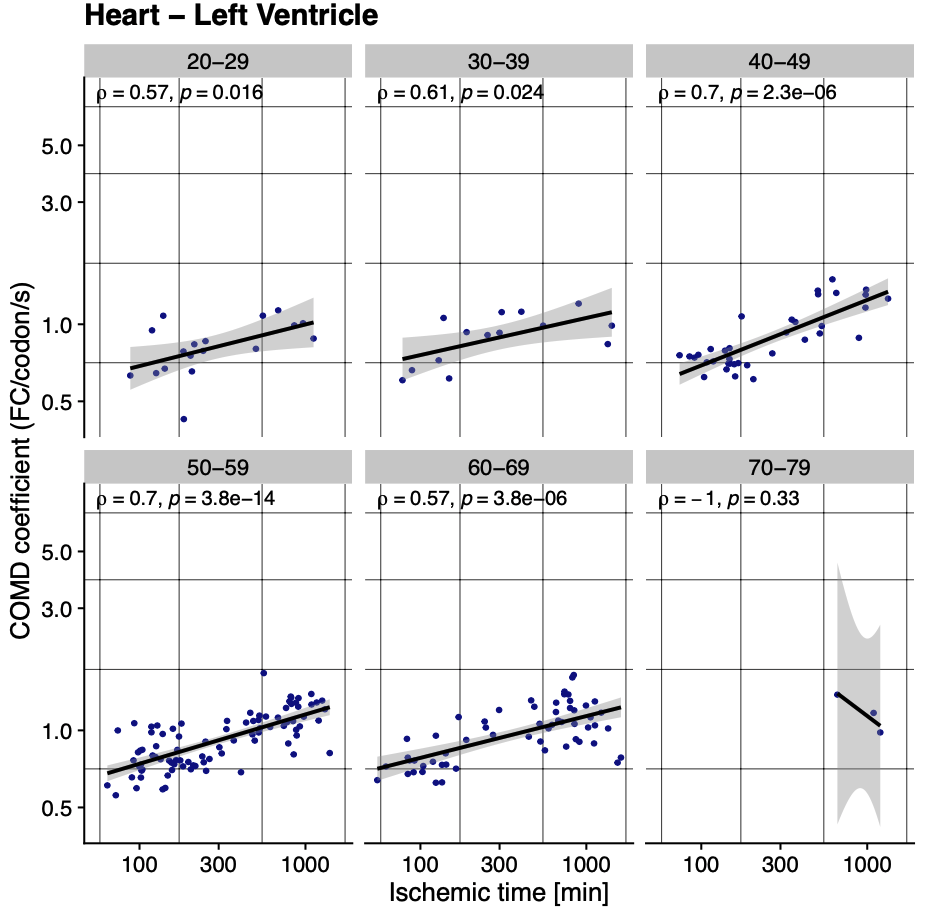
**Figure S7: The COMD coefficient associates with ischemic time for individuals of similar age in Heart - Left Ventricle.**  COMD coefficient against ischemic time (min) for Heart - Left Ventricle in different age groups.

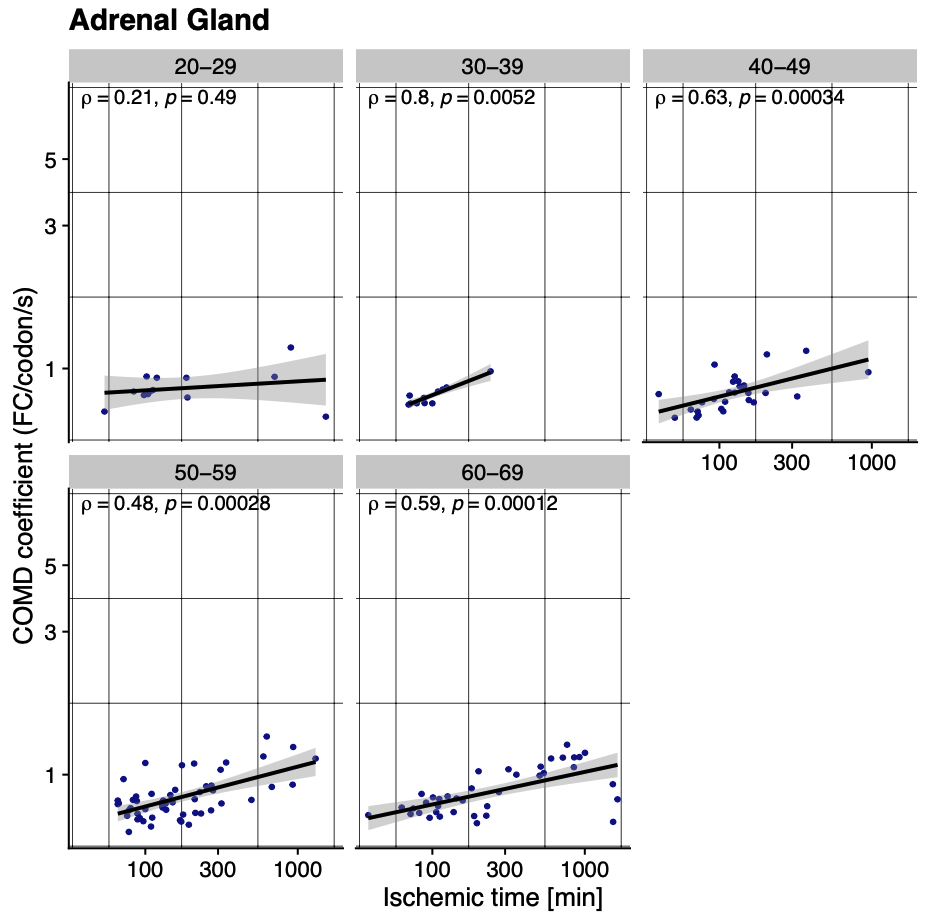
**Figure S8: The COMD coefficient associates with ischemic time for individuals of similar age in Adrenal Gland.**  COMD coefficient against ischemic time (min) for Adrenal Gland in different age groups.

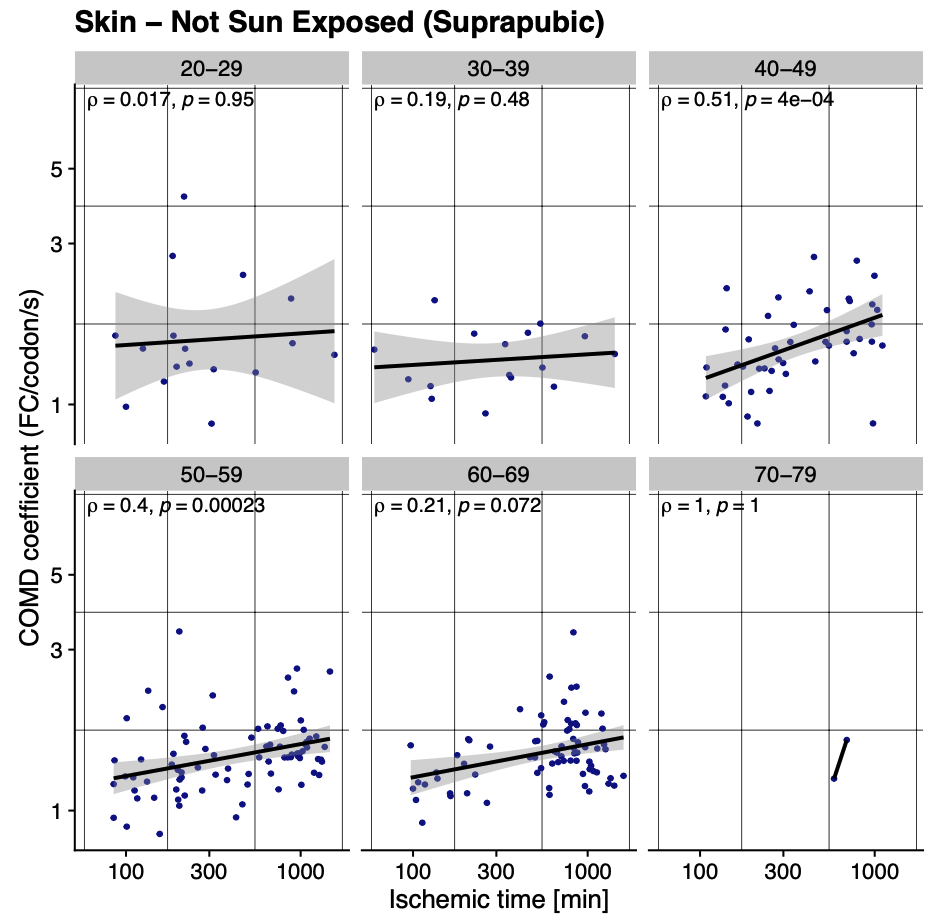
**Figure S9: The COMD coefficient associates with ischemic time for individuals of similar age in Skin - Not Sun Exposed (Suprapubic).**  COMD coefficient against ischemic time (min) for Skin - Not Sun Exposed (Suprapubic) in different age groups.

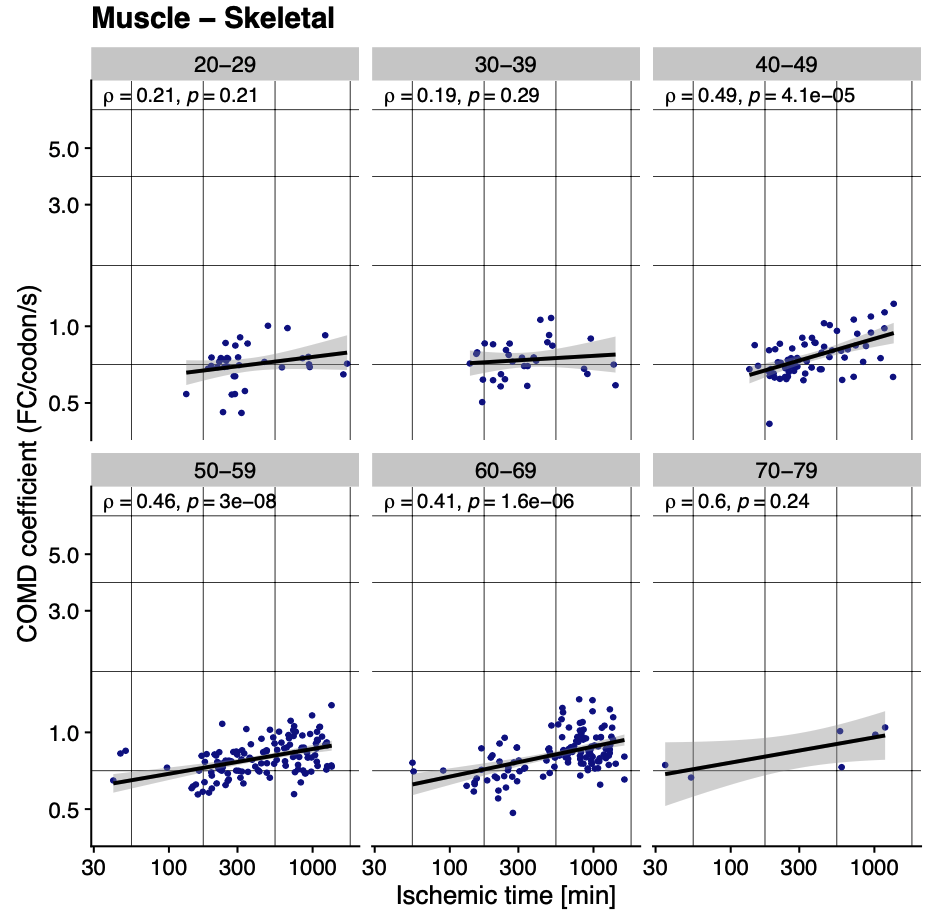
**Figure S10: The COMD coefficient associates with ischemic time for individuals of similar age in Muscle - Skeletal.**  COMD coefficient against ischemic time (min) for Muscle - Skeletal in different age groups.

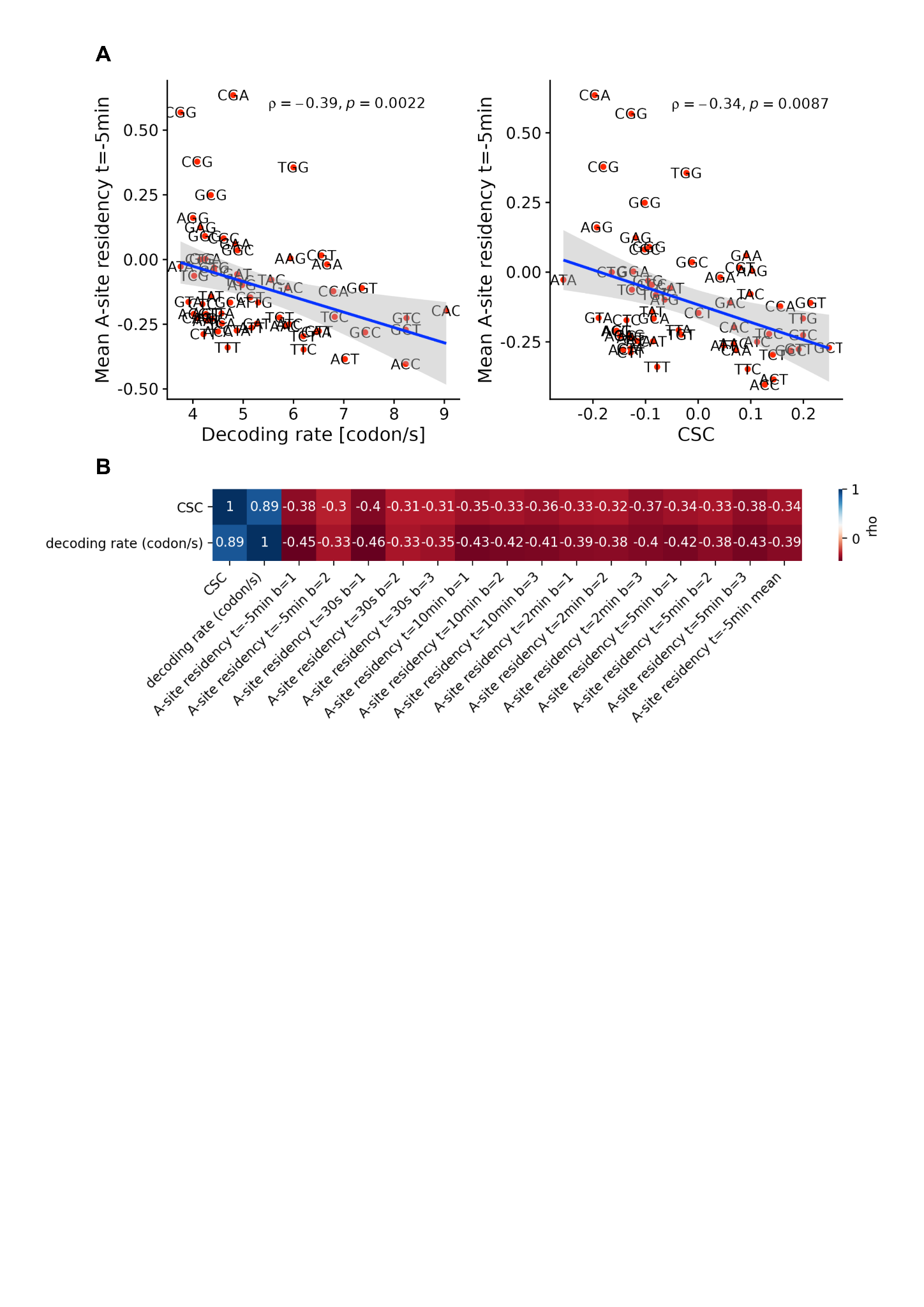

**Figure S11: Ribosome A-site residency estimated from 5Pseq correlates with codon optimality and decoding rate metrics. A** Mean A-site residency across batches at time point -5 min against CSC (codon stabilization coefficient, taken from [49]) and decoding rate (taken from [17]). **B** Correlation between ribosome A-site residencies of each sample (time point, *t*, and batch, *b*) and codon optimality or codon decoding rate.

| Term | Normalized Enrichment Score (NES) | False Discovery Rate (FDR) |
| --- | --- | --- |
| mitochondrial ATP synthesis coupled electron transport (GO:0042775) | -5.778 | <1.000E-06 |
| respiratory electron transport chain (GO:0022904) | -5.527 | <1.000E-06 |
| mitochondrial electron transport, NADH to ubiquinone (GO:0006120) | -5.256 | <1.000E-06 |
| mitochondrial electron transport, cytochrome c to oxygen (GO:0006123) | -4.049 | <1.000E-06 |
| mitochondrial respiratory chain complex I biogenesis (GO:0097031) | -3.927 | <1.000E-06 |
| mitochondrial respiratory chain complex I assembly (GO:0032981) | -3.927 | <1.000E-06 |
| NADH dehydrogenase complex assembly (GO:0010257) | -3.927 | <1.000E-06 |
| mitochondrial respiratory chain complex assembly (GO:0033108) | -3.893 | <1.000E-06 |
| pyruvate metabolic process (GO:0006090) | -3.841 | <1.000E-06 |
| mitochondrial ATP synthesis coupled proton transport (GO:0042776) | -3.840 | <1.000E-06 |
| ATP synthesis coupled proton transport (GO:0015986) | -3.583 | <1.000E-06 |
| cellular respiration (GO:0045333) | -3.219 | <1.000E-06 |
| mitochondrial translational termination (GO:0070126) | -3.184 | <1.000E-06 |
| ATP biosynthetic process (GO:0006754) | -3.089 | 7.511E-06 |
| cristae formation (GO:0042407) | -3.084 | 7.010E-06 |
| SRP-dependent cotranslational protein targeting to membrane (GO:0006614) | 2.987 | <1.000E-06 |
| cotranslational protein targeting to membrane (GO:0006613) | 2.921 | <1.000E-06 |
| ATP metabolic process (GO:0046034) | -2.903 | 6.572E-06 |
| nicotinamide nucleotide metabolic process (GO:0046496) | -2.877 | 6.185E-06 |
| mitochondrial translational elongation (GO:0070125) | -2.843 | 1.168E-05 |
| protein targeting to ER (GO:0045047) | 2.824 | <1.000E-06 |
| proton transport (GO:0015992) | -2.752 | 3.874E-05 |
| purine ribonucleoside triphosphate biosynthetic process (GO:0009206) | -2.717 | 5.783E-05 |
| cytoplasmic translation (GO:0002181) | 2.692 | <1.000E-06 |
| hydrogen ion transmembrane transport (GO:1902600) | -2.660 | 7.010E-05 |
| inner mitochondrial membrane organization (GO:0007007) | -2.612 | 9.081E-05 |
| viral gene expression (GO:0019080) | 2.603 | <1.000E-06 |
| translational termination (GO:0006415) | -2.564 | 1.463E-04 |
| viral transcription (GO:0019083) | 2.547 | <1.000E-06 |
| nuclear-transcribed mRNA catabolic process, nonsense-mediated decay (GO:0000184) | 2.520 | <1.000E-06 |
| aerobic respiration (GO:0009060) | -2.505 | 2.147E-04 |
| purine ribonucleoside monophosphate biosynthetic process (GO:0009168) | -2.307 | 1.039E-03 |
| peptide biosynthetic process (GO:0043043) | 2.287 | 1.340E-04 |
| canonical glycolysis (GO:0061621) | -2.100 | 4.210E-03 |
| glycolytic process through glucose-6-phosphate (GO:0061620) | -2.100 | 4.210E-03 |
| glucose catabolic process to pyruvate (GO:0061718) | -2.100 | 4.210E-03 |
| nuclear-transcribed mRNA catabolic process (GO:0000956) | 2.039 | 7.667E-03 |
| long-chain fatty acid transport (GO:0015909) | -2.019 | 7.335E-03 |
| negative regulation of cellular response to growth factor stimulus (GO:0090288) | 2.015 | 9.732E-03 |
| regulation of release of cytochrome c from mitochondria (GO:0090199) | -2.007 | 7.711E-03 |
| hexose biosynthetic process (GO:0019319) | -1.991 | 8.256E-03 |
| coenzyme biosynthetic process (GO:0009108) | -1.974 | 9.148E-03 |

**Table S1: Gene set enrichment analysis (GSEA) results for GTEx across tissues.**
GSEA was performed on the genes scored by their Spearman correlation between their expression (measured in TPM) and the COMD coefficient across tissues. Each gene set (row) is characterized by a GO term, a normalized enrichment score (NES) and a false discovery rate (FDR). Only gene sets with FDR < 0.01 were considered significantly enriched. Gene sets are ordered by their absolute value of the normalized enrichment score.
